# Design of Protein Sequences with Precisely Tuned Kinetic Properties

**DOI:** 10.1101/2025.02.13.638027

**Authors:** Z. Faidon Brotzakis, Michele Vendruscolo, Georgios Skretas

**Affiliations:** Institute for Bioinnovation, Biomedical Sciences Research Center “Alexander Fleming”, Flemingk 34, Vari, 16672, Greece; Centre for Misfolding Diseases, Yusuf Hamied Department of Chemistry, University of Cambridge, 12 Union Rd, Cambridge, CB2 1EZ, United Kingdom

**Keywords:** Inverse Folding, Molecular Dynamics, Protein Design, Enhanced Sampling Molecular Dynamics, Metadynamics, Protein Dynamics, Antibody design, SARS-CoV-2, Molecular Kinetics, Bayesian Optimization

## Abstract

Recent advances in computational biology have enabled solutions to the inverse folding problem - finding an amino acid sequence that folds into a target structure. An open question concerns the design of proteins that in addition to having the correct target structure also have precisely tuned kinetic properties, such as folding and unfolding rates. To address this problem, we formulate the inverse folding problem as a quest for a sequence with a target free energy landscape. To propose a procedure to address this problem, here we describe the Inverse Folding Molecular Dynamics (IF-MD) method, which combines inverse folding with enhanced sampling molecular dynamics and Bayesian optimization. IF-MD leverages ensemble averages from molecular dynamics simulations, reweighted according to a Bayesian framework, to guide the design of sequences exhibiting specific kinetic properties. We demonstrate the methodology by optisising the binding kinetics of H11, a nanobody against the SARS-CoV-2 spike receptor-binding domain (RBD), thus identifying nanobody variants with slower unbinding kinetics than H11. Mechanistic analysis reveals that this kinetic property arises from a shift towards configurations closer to the bound state and increased free energy barriers for dissociation. These findings highlight the power of IF-MD for efficiently navigating the vast sequence space to design proteins with a tailored free energy landscape.

## Introduction

Advances in computational methods for the inverse folding problem, in particular using deep-learning, have enabled the design of protein sequences with predefined native states, and capable of forming stable complexes with chosen partners [1–4].

As conformational dynamics play a crucial role in protein function and regulation [5], an open problem concerns the design of kinetic properties [6, 7]. Developments in this direction will improve the accurate prediction of protein behavior in complex biological systems.

The design of proteins with specific functionalities and dynamic properties is a central challenge in various fields, including biotechnology, biomedicine, and materials science [8, 9]. Traditional approaches to protein design often rely on iterative cycles of design, simulation, and experimental validation, which can be computationally expensive and time-consuming [10]. Furthermore, accurately predicting and controlling the kinetics of protein folding and binding remains a significant hurdle, especially for rare events that occur on slow timescales relevant to many biological processes [11].

This work introduces a computational approach to efficiently explore simultaneously the sequence space and the conformational space of proteins, and thus design proteins with precisely tuned kinetic properties. Our strategy combines three key elements: inverse folding molecular dynamics (IF-MD), Bayesian optimization, and enhanced sampling. First, IF-MD leverages ensemble averages from molecular dynamics (MD) simulations, reweighted according to a Bayesian framework [12], to guide the design process. These ensemble averages, which represent macroscopic observables such as binding affinity or unbinding rate constants, act as constraints that direct the search for optimal protein sequences. Second, Bayesian optimization, a global optimization technique [13], efficiently navigates the vast sequence space and proposes new sequences that are likely to exhibit the desired macroscopic properties. Finally, to enhance sampling efficiency and overcome kinetic barriers associated with slow conformational transitions, the IF-MD framework is coupled with infrequent metadynamics, a powerful enhanced sampling method that enables exploration of complex free energy landscapes to predict transition rates[14, 15]. This combination of techniques enables fast sampling of relevant conformational states, ensuring that the Bayesian optimization procedure is provided with sufficient statistical data for robust predictions and leading to more efficient and accurate protein designs.

Starting from a known low-affinity lead nanobody binder to SARS-CoV-2 RBD derived from phage display [16], we demonstrate this methodology by applying it to the design of nanobodies with precisely tuned slow unbinding kinetics from the SARS-CoV-2 spike receptor-binding domain (RBD).

## Results

### Inverse Folding Molecular Dynamics

Free energy landscapes enable the representation of the structure, thermodynamics and kinetics of proteins [17] (Fig. 1A). MD simulations provide a tool to generate free energy landscapes and to characterise folding and unfolding events, as well as binding and unbinding rates.

**Fig. 1.**
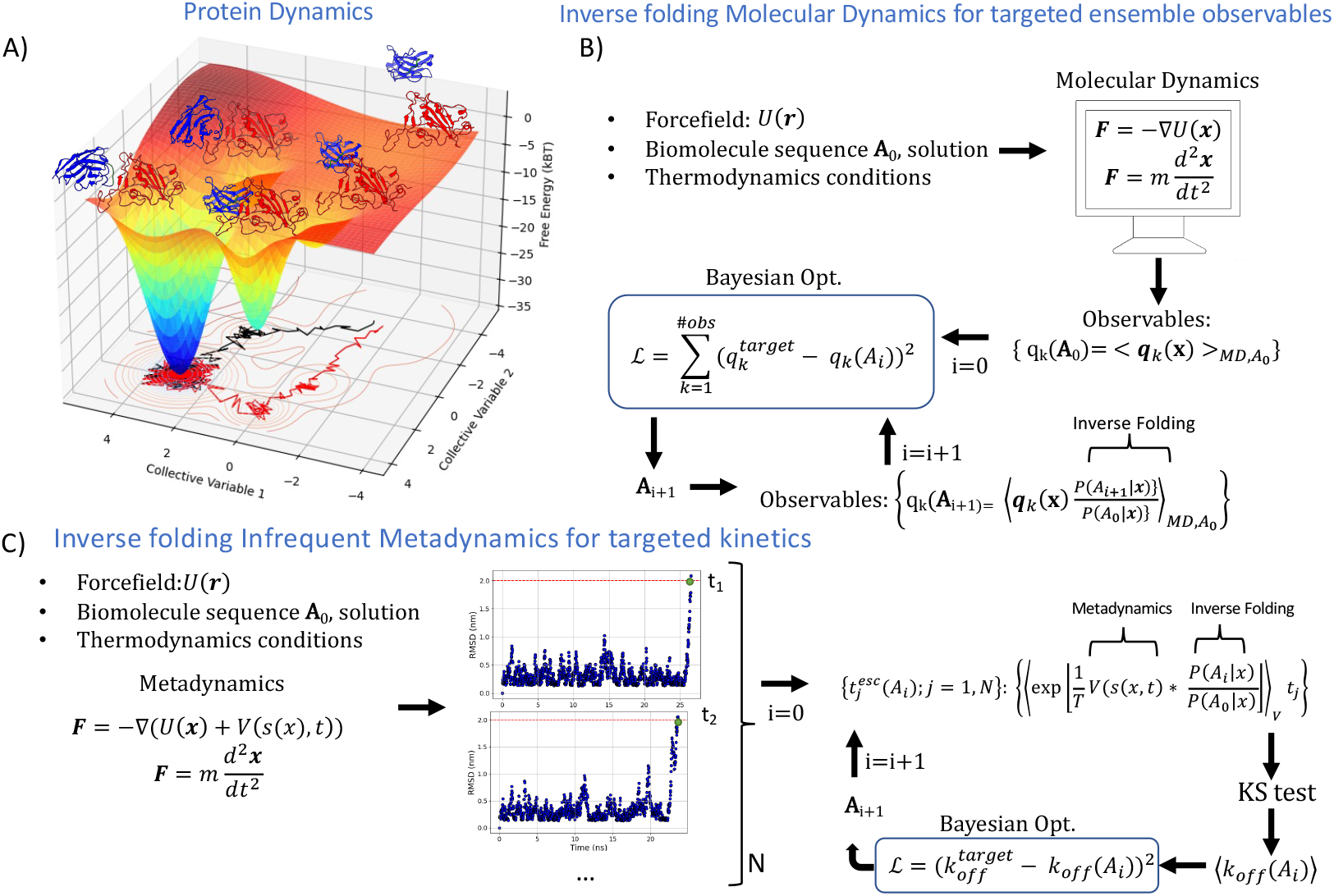
Inverse Folding Molecular Dynamics (IF-MD) protocol. (A) Free energy landscape of the process of protein-protein dissociation in the case of SARS-CoV-2 RBD (red) in complex with the nanobody H11 (blue) visiting different metastable states along the unbinding mechanism along two collective variables. Two different unbinding trajectories from the bound state are shown in black and red. (B) IF-MD protocol to sample protein sequence space by constraining *k* ensemble averaged observables 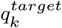 and using Bayesian Optimization to propose new sequences. (C) Specific example of IF-MD constrained to a target the unbinding rate constant, and using Infrequent Metadynamics to sample the prior ensemble. Unbinding example trajectories sampled by Infrequent Metadynamics are shon in blue, along the RMSD from the cryo-EM structure of H11 bound to SARS-CoV-2 RBD as a function of time.

Inverse Folding Molecular Dynamics (IF-MD) provides a scheme to navigate the protein sequence space to optimize target macroscopic observables based on ensemble averages from prior MD trajectories. In IF-MD, these MD trajectories are reweighted by an inverse folding scheme (e.g. MPNN [12], see Methods) and by utilizing Bayesian optimization equipped with a loss functions related to the target macroscopic property of interest (see Fig. 1B and Methods). To obtain long timescale dynamics, IF-MD can be coupled to enhanced sampling MD methods [18–20]. In this study, we combine IF-MD with Infrequent Metadynamics [14] to enable fast sampling of the distribution of nanobody/RBD unbinding pathways used to predict corresponding unbinding rate constants and design nanobody sequences with targeted unbinding kinetics.

### Anti RBD nanobody optimization for targeted slow unbinding kinetics

Starting from Inf-MetaD simulations of an RBD/H11 nanobody complex (PDB:7Z6V) exhibiting a mean unbinding escape time of (1.02 10^−5^ s), we applied IF-MD to iteratively design nanobody variants by altering the CDR3 sequence of residues 100-104 (20^5^ sequence space) and targeting a 10^−3^ s unbinding escape (Fig. 2A). We found that this constraint optimization proposes nanobody sequences with slower unbinding kinetics than H11 up to two orders of magnitude. Notably, we selected nanobody designs N and R as negative controls as they appeared to show faster unbinding kinetics than H11, while D, A and AR exhibited slowed unbinding kinetics (Fig. 2B and Table 1). We rationalized the IF-MD derived change in kinetics by plotting in Fig. 2C the probability distribution projection along RMSD from the cryo-EM H11/RBD structure of the different IF-MD ensembles. Nanobodies with slowed unbinding kinetics exhibited higher densities close to the bound state, consistently with the slower unbinding escape times. To validate the accuracy of IF-MD, we performed validation simulations in Inf-MetaD where we identified the unbinding kinetics of all designed nanobody variants and found a correlation between the IF-MD and the validation Inf-MetaD derived unbinding escape times (Fig. 2D). However it appears that the agreement between the two increases when at IF-MD p-values higher than 0.15 (Fig. 2D,inset and Table 1). Therefore we deemed the p-value to be a sensitivity criterion of the accuracy of the IF-MD predictions. Nearly all nanobody designs (Fig. 2E and Table 1) showed slower unbinding kinetics from the RBD than H11 and maintain the original nanobody fold as quantified by the low RMSD of the designs with respect to H11 by using the NANONET predictor [21] (Table 1). The increased potency of all variants compared to H11 shown in the validation step is further verified using Rosetta[22] molecular docking calculations, by the higher affinity and greater number of contacts formed between the nanobody CDRs and the RBD epitope.

**Table 1.** Statistics of escape times and associated p-value and k_*off*_ for each system in the IF-MD(Designs) and reruns (Validaiton).RMSD comparison of structure of nanobodies with respect to H11.

| Designs |  |  |  |  |
| --- | --- | --- | --- | --- |
| Nanobody System | Escape time (s) | $k_{off}(1/s)$ | p-value | $RMSD(\text{\AA}; H11)$ |
| N | 1.45E-06 | 6.90E+05 | 1.6E-01 | 0.11 |
| R | 5.75E-06 | 1.74E+05 | 1.5E-01 | 0.12 |
| H11 (Mikolajek et. al.) | 1.02E-05 | 9.80E+04 | 1.89E-01 | 0 |
| D | 1.97E-04 | 5.08E+03 | 1.3E-01 | 0.11 |
| A | 3.53E-04 | 2.83E+03 | 1.4E-01 | 0.12 |
| AR | 8.39E-04 | 1.19E+03 | 1.73E-01 | 0.11 |
| H11-D4 (Mikolajek et. al.) | 1.40E-04 | 7.15E+03 | 1.5E-01 | 0.11 |
| Validation |  |  |  |  |
| N | 7.52E-05 | 1.33E+04 | 7.62E-02 |  |
| R | 1.23E-05 | 8.16E+04 | 1.20E-01 |  |
| D | 3.72E-05 | 2.69E+04 | 1.80E-02 |  |
| A | 4.70E-05 | 2.13E+04 | 2.31E-02 |  |
| AR | 2.13E-04 | 4.70E+03 | 1.38E-01 |  |
| H11-D4 (Mikolajek et. al.) | 1.05E-04 | 9.49E+03 | 0.1 |  |

**Fig. 2.**
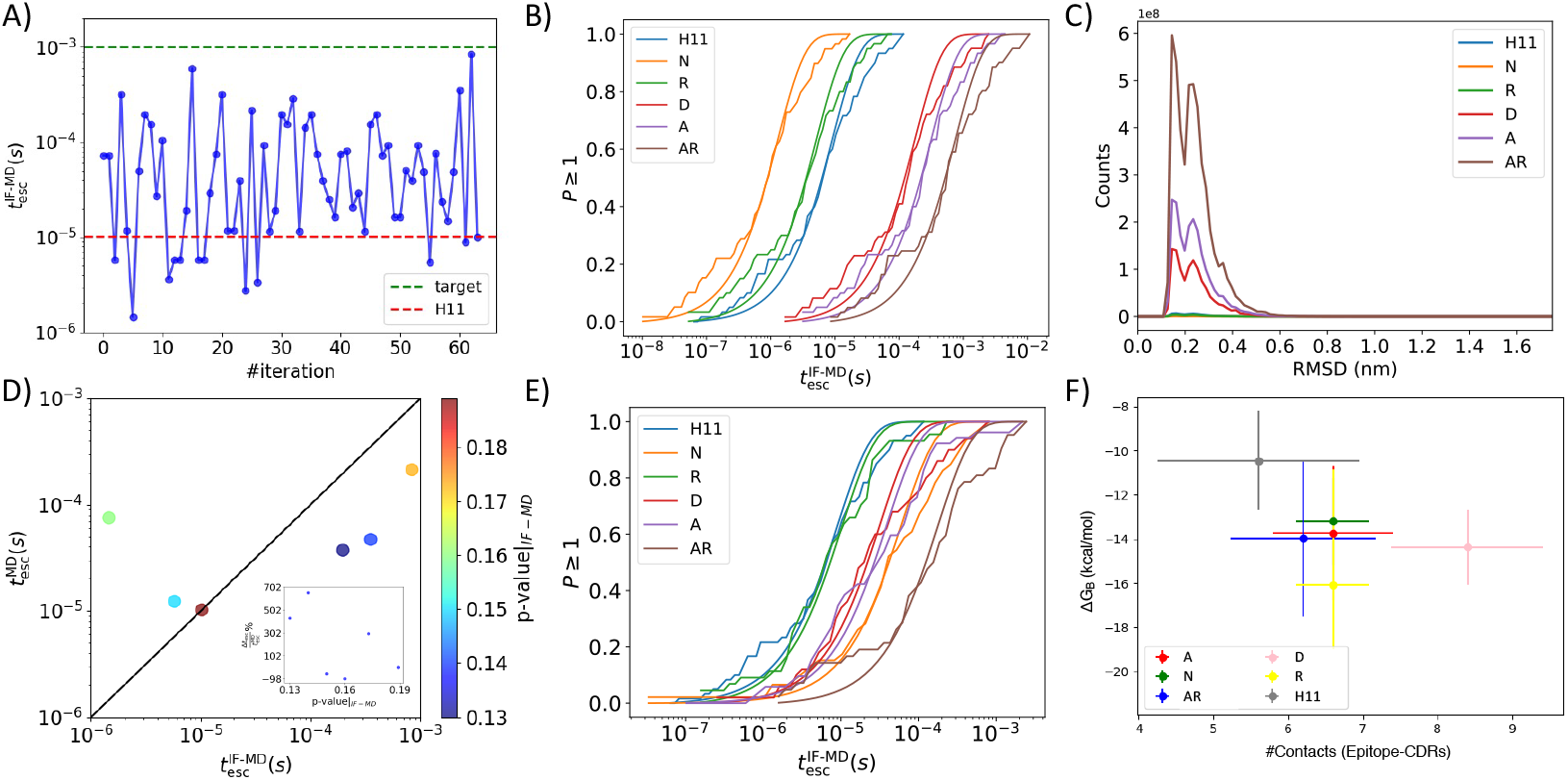
Nanobody optimization for targeted RBD/H11 unbinding kinetics. (A) Inverse-Folding unbinding escape time as a function of iteration. The H11/RBD unbinding escape time is shown in red/black. (B) Empirical (thin lines) and theoretical CDF of unbinding escape times of designed nanobodies either directly evaluated from the IF-MD simulations (thin lines) and fitted as a Poisson process (solid lines). P_*n*≥1_ denotes the probability of observing *n* events. (C) Probability distribution projection along RMSD from cryo-EM H11/RBD of the different IF-MD ensembles. (D) Unbinding escape times of designed nanobodies from RBD obtained from IF-MD versus validation Inf-MetaD simulations with p-value of IF-MD escape time statistics in heat map. The inset shows the relative error of IF-MD unbinding escape times versus p-value of IF-MD. (E) Empirical (thin lines) and theoretical CDF of escape times either directly evaluated from the validation MD simulations (thin lines) and fitted as a Poisson process (solid lines). P_*n*≥1_ denotes the probability of observing *n* events. (F) Molecular docking predictions of the binding energy of designed nanobodies as a function of nanobody CDR1 (28-32),CDR2 (55-56),CDR3 (101-107) and the RBD epitope PRO491-SER494.

### Unbinding mechanism of nanobodies from RBD

We compared the AR nanobody variant, which showed 20-fold slower unbinding kinetics from RBD, to H11 in the validation Inf-MetaD, in mechanistic free energy landscape analysis. By comparing the H11 IF-MD landscapes (Fig. 3A,D,G) to the AR ones (Fig. 3B,E,H), we observed a common trend of decreasing the population of configurations outside of the ground state and along CVs distance (*d*_102−494_ ≥ 1.5nm) and RMSD (at RMSD ≥ 1nm), in par with increasing the free energy barrier for dissociation. In the validation Inf-MetaD simulation of AR, we distinguished two mechanisms along unbinding shown in Fig. 3C,F,I and in the appendix Fig. 1A-C. First, a sliding mechanism, where the protein reorients along the surface to an intermediate state B1, rebinds to the ground state B0 before dissociating to the unbound state U, and second an aligned mechanism where the nanobody directly unbinds towards the unbound state.

**Fig. 3.**
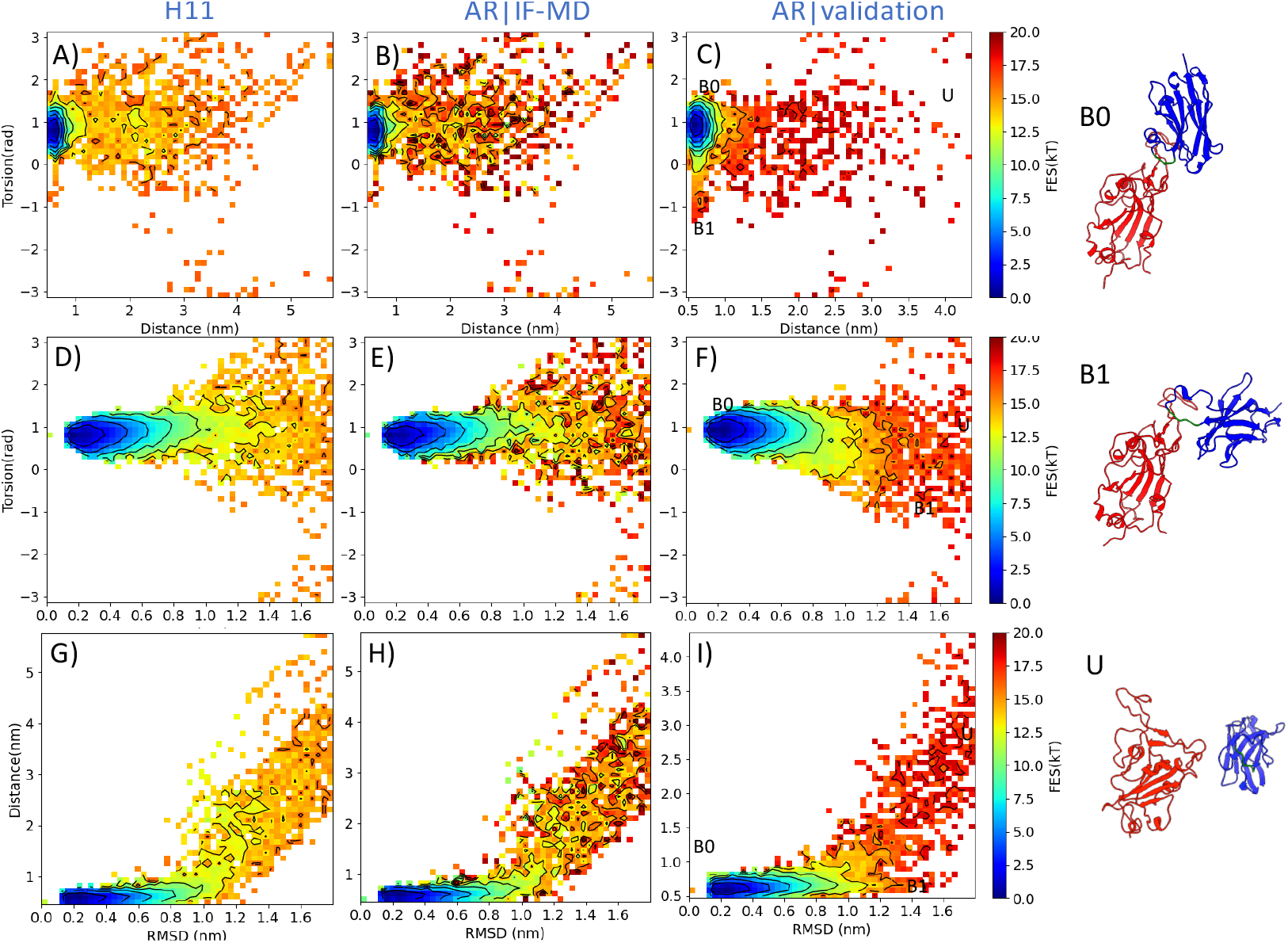
Unbinding mechanism of nanobodies from RBD. Free energy surface projection along distance d_102−494_ and torsion descriptor *ϕ* (A-C), torsion descriptor *ϕ* and RMSD from ground state complex (D-F) and distance d_102−494_ and RMSD form ground state complex (G-I) for H11-RBD prior simulation, AR-RBD IF-MD and AR-RBD Inf-MetaD validation respectively. Metastable states identified in the AR-RBD Inf-MetaD validation simulation denoted as B0 (ground state), B1 (intermediate) and U (unbound) are found on the panel on the right.

## Discussion

We reported the IF-MD method to include protein dynamics in inverse folding procedures, thus enabling an estimate of the free energy of binding not just based on energy, but also on entropy. The MD engine, force field and the enhanced sampling scheme are user-specific, while in the optimization scheme the user can incorporate tailored observable terms in the loss function, or change the optimizer to other black box optimization algorithms with feedback such as reinforcement learning [13, 23] and genetic algorithms [24]. IF-MD produces fast estimations of probability distributions in the conformational space, and respective thermodynamic and kinetics macroscopic observables by using the reweighting in eqs. 3,5, thereby assuming a similar mechanism of action of the prior ensemble with the reweighted ones for different protein sequences. We note that as in other reweighting approaches, this procedure becomes increasingly accurate as a wider prior ensemble is explored [25, 26]. In this study we showed that the higher the p-value of nanobody-RBD unbinding escape time statistics of the IF-MD reweighted ensemble of a new nanobody sequence is, the higher the agreement with the validation MD unbinding escape time estimate, hence p-value IF-MD can be a sensitivity parameter of how close to the original mechanism we are.

To illustrate the IF-MD method, we described the AR nanobody, which showed slower unbinding escape times from RBD compared to the parent H11 nanobody. This behavior is generated by the presence of two distinct unbinding mechanisms, a sliding and an aligned one, as opposed to H11, which shows only the latter. Such mechanisms were previously identified in protein-protein dissociation and association [15, 27, 28]. Quite generally, low populated metastable states often act as kinetic traps [29–31], hindering the system from reaching its global minimum efficiently. The system may spend a prolonged time in these trapped conformations before successfully overcoming free energy barriers to transition to a more stable state. This slows down the overall binding process. In the case studied here, the frustrated intermediate metastable states B1, occurring along the sliding unbinding mechanism, acts as a kinetic trap by increasing the rebinding probability to the ground state, thus decreasing the unbinding rate and increasing the stability of the bound complex. More generally, the presence of frustrated free energy landscapes can diversify the possible pathways that a protein can take to reach its native or final bound state. IF-MD offers the possibility of engineering the free energy landscape, in addition to the ground state structure, thus making it possible to tailor the thermodynamic and kinetic properties of the designed sequences.

## Methods

### Protein Design Protocol

In protein design, or inverse folding, the quest is to find amino acid sequences able to adopt a predetermined structure [2, 12]. In other words, one searchers for protein sequences that maximize *P* (sequence | structure), the Boltzmann probability that a sequence occupies a given structure [32]. Widely used deep learning inverse folding models such as MPNN [12] and ESM-IF [2] approximate this conditional distribution and enable the relative comparison of different sequences to adopt a predetermined fold.

### Reweighting algorithm

Consider a protein-binder comprised of *N* residues, with sequence *A* = {*A*_1_, …, *A*_*N*_} and a target of *M* residues *T* = {*T*_1_, …, *T*_*M*_}, where residues can span the 20 natural amino acids space and *X*, the Cartesian atomic coordinates of the complex. Suppose that we want to compare two protein sequences, e.g protein-binders, *A* and *Â* for their relative likelihood to form a structural complex with the target, *X*. By using Bayes’ formula, we can compare two sequences *A* and *Â* as:

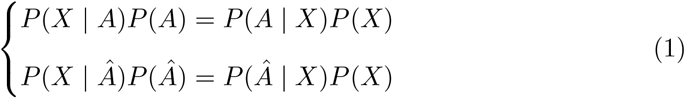

where *P* (*X*) is the probability of a structure, *P* (*A*) and *P* (*Â*) are the probabilities of sequences. After rearranging, eq. 1 becomes:

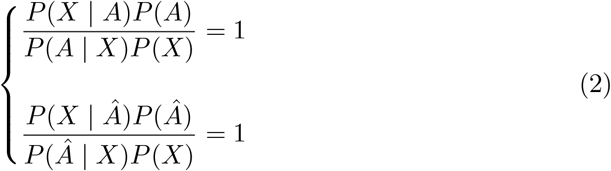

After equating the right-hand side of the two sets of equations in eq.2, rearranging and considering cancellation of the *P* (*x*) terms, we obtain:

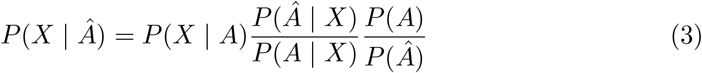

In the right-hand side of eq. 3, the first term denotes a prior probability distribution of configurations X given a sequence *A* that can be predicted by MD generating configurations of the protein binder sequence *A* that we wish to redesign, the target *T* and the environment (e.g water solution, membrane, ion, co-solutes, pH) depending on the context of the target under study. The second term by Protein MPNN or ESM, and the third by MSA-based Potts models [33–35]. The aforementioned formalism assumes that the configurational dynamics of the prior *P* (*X* | *A*) is similar to *P*(*X* | *Â*) and it just accounts to reweight by the second and third factors of the right-hand side of eq. 3.

### Loss function and optimization

To navigate the sequence space of the binder in order to match a target value 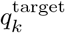 of an ensemble averaged property *k, q*_*k*_(*A*) = ⟨*q*_*k*_(*X*)⟩_MD,*A*_ we define the following loss function.

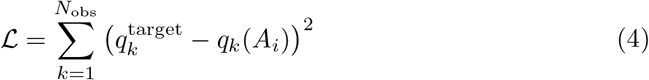

Starting from a molecular dynamics trajectories giving access to ensemble averaged properties at iteration *i* = 0, *q*_*k*_(*A*_0_) = ⟨*q*_*k*_(*X*)⟩_MD,*A*0_, (in this study *k*_*off*_, *RMSD*(*Â*; *A*_0_)),and defining target values of 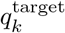 (in this study 10^−3^ s and 0.1 nm), the loss in 4 is calculated and using a Bayesian optimization based on Gaussian process regression implemented in the Python gp minimize function, a new sequence *Â* = *A*_*i*+*i*_ is proposed. Then we perform reweighting by using eq. 3 and approximating *P* (*Â*) as *P* (*Â*) ∝ exp [*RMSD*(*Â*; *A*_0_)], where NANONET [21] are used to predict the fold of sequence *Â*. Since we determined in the loss function that nanobody sequences sample from the original nanobody family, that is they fold to very similar structures to the parent H11 nanobody (*A*_0_) fold (*RMSD*(*Â*; *A*_0_) *<* 0.1(nm)) (see Table 1), terms *P* (*Â*) and *P* (*A*) are approximately equal and can be neglected. Hence at each iteration *i*, the observable *k* can be written as

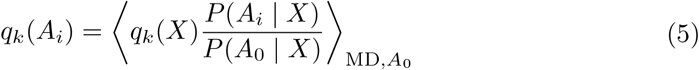

where structures appear according to the MD distribution e.g. Boltzmann at NVT, and the second right hand side term can be calculated using Protein MPNN. The pair (*A*_*i*=*i*+*i*_, *q*_*k*_(*A*_*i*=*i*+1_)) enters in the next step in eq.4. The protocol is summarized in Fig.1B, and for the specific case of targeting unbinding rate constants by using Infrequent Metadynamics to sample the prior ensemble *P* (*X* | *A*_0_) following Baysian Optimization of sequences, in Fig.1C.

### Nanobody unbinding rates from Infrequent-Metadynamics Molecular Dynamics

To tackle the rare event problem of MD and enable reaching ligand-target dissociation events on the ms-s timescale, typically occurring in antibody-antigen recognition, we used Infrequent-Metadynamics [14, 36]. Inf-Metad deposits bias along a collective variable at high pace and its accuracy in rate estimate prediction relies on: (i) no bias deposited at the transition state ensemble (TSE) region, and (ii) ability of biased CVs to distinguish between relevant metastable states. These conditions are met by employing infrequent bias deposition, thus minimizing the probability of adding bias to the short transition path time where the system lies at the TSE.

Metadynamics accelerates the conformational dynamics by a factor *α*, calculated by a running average obtained in the Metadynamics trajectory as: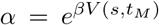, where *t*_*M*_ is the metadynamics escape time, *s* is the biased collective variable at time *t*_*M*_, *β* is the inverse of *k*_*B*_*T*, and *V* (*s, t*_*M*_) is the bias deposited at time *t*_*M*_. The real escape time *t* can be related via:

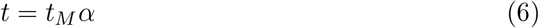

A rigorous quantitative way to assess the assumption of the method is to perform a Kolmogorov–Smirnov statistical test[37]. This effectively measures if the *M* observed escape times follow a Poisson distribution, as expected in a rare event scenario.

### Kolmogorov-Smirnov Test

The nanobody-RBD unbinding transition is a rare event, and the likelihood of observing an individual transition at time *τ* is distributed exponentially, characterising a homogeneous Poisson processes. To rigorously assess whether the process is Poissonian and therefore no bias has been deposited on the TSE region, in the Kolmogorov-Smirnov test we compare the empirical cumulative distribution function (ECDF) of N individual escape times,each obtained from eq. 6, with a theoretical cumulative distribution function (TCDF), which for homogeneous Poisson process is defined as TCDF=1-exp(-t/*τ*). We use the p-value of the KS statistics as a measure of similarity between the TCDF and ECDF and actually quantifies the likelihood that the escape times by Inf-MetaD follow the rare event theoretical exponential distribution. A trustworthy estimate of the observed escape times in Inf-MetaD requires,the p-value, to be greater than the confidence level of 0.05. The transition considered in this manuscript is the full unbinding transition, that is the time it takes for the ligand to escape the bound state and land in the unbound state (RMSD*>*1.8 nm). The k_*off*_ is considered as the reciprocal of the estimator of the TCDF.

### Infrequent Metadynamics Simulation Details

#### H11 simulations

We started from the cryo-EM based structure of H11 bound to RBD identified in a previous study[16] (PDB: 7Z6V) and kept a single RBD-H11 complex that lying in the up conformation (chain A), spanning RBD residues ASN334-LYS529 and the whole H11 nanobody (chain D). We then used SMOG2[38] to construct a contact potential force field following exactly the same GROMACS MD parameters and setup steps as in[39], which are summarized as: (a) construct contact potential, (b) energy minimization, (c) MD equilibration in the NVT for 2 ns at 298 K in GROMACS 2020.5 [40], and (d) carry on 60 Inf-MetaD simulations starting with different velocities from the last frame of the previous step, with upper simulation time of 100 ns, and stopping conditions if RMSD*>*1.8 nm, where the nanobody was considered dissociated. For all cases, as collective variable we considered the RMSD from the cryo-EM structure (PDB: 7Z6V). Gaussians were deposited every 40 ps, a well-tempered Metadynamics bias factor of 7 and a Gaussian height of 2.5 kJ/mol has been used. The Gaussian height and bias factor is shown in Table S1 per system. We also keep track of two additional descriptors, i.e the distance between the C*α* of the central residue of CDR3 center (100RVTRS104), R102 and the center of the RBD epitope, SER494, abbreviated as d_102−494_. Moreover, we kept track of a dihedral descriptor *ϕ* highlighting the orientation of the complex comprising N atoms of RBD ASN354, VAL510 and H11 ALA97, ALA50. We added upper RMSD walls to 0.1 for each of RBD or nanobody to prevent it from losing tertiary structure upon biasing the total RMSD.

#### Validation of nanobody designs simulations

We followed the same protocol described above to run the Infrequent Metadynamics of the validation simulations of the nanobody designs from the molecular dynamics inverse folding simulations. The starting structure upon which the SMOG2 contact potential is made in step (a) is the most probable state occurring in the inverse folding reweighted ensemble.

#### Molecular Docking Validation

For each nanobody design, we utilized the Rosetta [22] docking software equipped with the Ref2015 scoring function [22] and performed 50 cycles of rigid docking and refinement, leading to 15 refined output structures, while using a harmonic constraint at d_102−494_ to 0.3 nm. For analysis, we quantified average and standard deviation between the 5 structures forming highest number of *Cα* contacts between between RBD PRO491-SER494 and any of CDR1 (28-32),CDR2 (55-56),CDR3 (101-107) as well as having the smallest interfacial binding energy.

## Acknowledgments

Z.F.B and G.S would like to acknowledge support from Horizon Europe Programme under the “Widening Participation Spreading Excellence” component (call ERA Chairs “HORIZONWIDERA-2022-TALENTS-01-01–ERA Chairs”); Project “Boost4Bio”; Grant Agreement No. 101087471. Z.F.B would like to acknowledge support from the National Infrastructures for Research and Technology (GRNET) for computer resources provided through the Amazon AWS project “PMDrescue”.

## Supplementary information

Additional information referenced in the text is available in the Appendix.

## Appendix

**Fig. 1.**
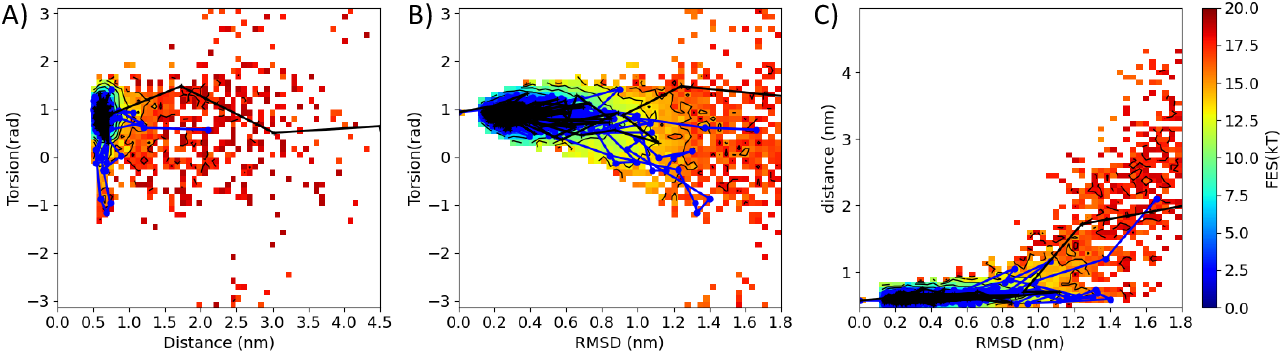
Unbinding Pathways. Free energy surface of AR-RBD validation Inf-MetaD simulations projected on A)d_102−494_-*ϕ*, B) RMSD-*ϕ*, C) RMSD-d_102−494_. With blue a characteristic sliding mechanism pathway and in black and aligned dissociation pathway.

